# Natural diversity of honey bee (*Apis mellifera*) gut bacteriome in various climatic and seasonal states

**DOI:** 10.1101/2021.01.27.428438

**Authors:** Márton Papp, László Békési, Róbert Farkas, László Makrai, Gergely Maróti, Dóra Tőzsér, Norbert Solymosi

**Affiliations:** University of Veterinary Medicine Budapest, Centre for Bioinformatics, Budapest, 1078, Hungary; University of Veterinary Medicine Budapest, Department of Parasitology and Zoology, Budapest, 1078, Hungary; University of Veterinary Medicine Budapest, Department of Microbiology and Infectious Diseases, Budapest, 1143, Hungary; Plant Biology Institute of the Biological Research Center, 6726 Szeged, Hungary; University of Public Service, Faculty of Water Sciences, 6500 Baja, Hungary; University of Veterinary Medicine Budapest, Department of Food Hygiene, Budapest, 1078, Hungary; Eötvös Loránd University, Department of Phyisics of Complex Systems, Budapest, 1117, Hungary

## Abstract

As pollinators and producers of numerous human consumed products, honey bees have great ecological, economic and health importance. The composition of their bacteriota, for which the available knowledge is limited, is essential for their body’s functioning. Based on our survey, we performed a metagenomic analysis of samples collected by repeated sampling. We used geolocations that represent the climatic types of the study area over two nutritionally extreme periods (March and May) of the collection season. In bacteriome composition, significant (p=0.002) difference was found between the samples from March and May. The samples’ bacteriome from March showed a significant (p=0.02) composition difference between cooler and the warmer regions. However, there were no significant bacteriome composition differences among the climatic classes of samples taken in May. Based on our results, one may conclude that the composition of healthy core bacteriome in honey bees varies depending on the climatic and seasonal conditions. This is likely due to climatic factors and vegetation states determining the availability and nutrient content of flowering plants. The results of our study prove that in order to gain a thorough understanding of a microbiome’s natural diversity, we need to obtain the necessary information from extreme ranges within the host’s health state.

## Introduction

Honey bees are important pollinators with high economic value and ecosystem importance^1–3^. Their economic significance is based on their role in crop pollination and the different bee products they make^2, 3^. Honey, their most well-known product, is an important component of the human diet. Some evidence suggests that honey consumption can improve human health and might have a role in disease management^4^. However, honey bees are exposed to confined environments, and several factors threaten their health, including different pathogens, parasites and chemicals used as pesticides in agriculture^5–7^. The global decline of this key pollinator poses a threat to food security and to the maintenance of biodiversity^8^. There is growing attention on the effects of different herbicides and pathogens on the bee gut microbiota^9–11^. However, there is no detailed evidence about the natural variability of the honey bee gut bacteriota. Nevertheless, this could form the basis of studies exploring the effect of different harmful agents on honey bees’ gut bacteriota. Without this knowledge, one cannot decide if any suspected factor places the bacteriota composition into an adverse state. Although there are studies on honey bee gut microbiota and microbiomes^12–14^, none were designed to describe the effect of feed on its composition. It is known for many other animals that feed affects the composite of gut bacteriota. Subsequently, it can be assumed that bee gut bacteria may also be altered by feed, which in turn can be affected by seasonal and environmental conditions. During the collection season, various flowering plants provide diverse feed for the bees. The vegetation cycles, flowering, and pollen quantity and quality of plants are mostly influenced by meteorological conditions, especially precipitation and temperature. Our study aimed to get a deeper insight into the natural variation of gut bacteriomes in healthy worker honey bees based on a country-wide, repeated measure survey. To achieve the research goal, we were guided by considering that it is advisable to work based on samples representing extreme states of bee gut bacteriota. Hence, samples were taken during the two most distinctive periods of the honey collection season and sampling sites were selected from markedly distinct areas based on their climatic characteristics.

## Materials and Methods

### Sampling design and sample collection

The study’s main goal was to understand the natural variability in the gut bacteriome of healthy honey bees (*Apis mellifera*) and we have designed the sampling to be representative of Hungary. For this purpose, a feasible way was to identify sampling points representative of climatic conditions. In Hungary, there are 175 local administrative units (LAU 1), for each of these units, we have calculated the 10-year average of the yearly growing degree days (GDD) with base 10°C^15, 16^ and the yearly total precipitation. Meteorology data for period 2008-2017 was gathered from the ERA-Interim reanalysis data repository^17^ by the spatial resolution of 0.125°. For both environmental variables, two-two categories were defined: cooler-warmer and less-more for GDD and precipitation, respectively. Regarding GDD the lower two quartiles were classified as cooler and the upper two quartiles as warmer. For precipitation, the yearly mean below the country-wide median was assumed as less, above the median as more. Each LAU 1 was categorised by its own climatic variables (Fig 1). By stratified spatial random sampling^18, 19^, twenty local administrative units were chosen as sampling areas. The strata’s sample size was proportional to the stratifying GDD and precipitation categories’ country-wide frequency in order to be representative. All data management and analysis were performed in the R environment^20^.

**Figure 1.**
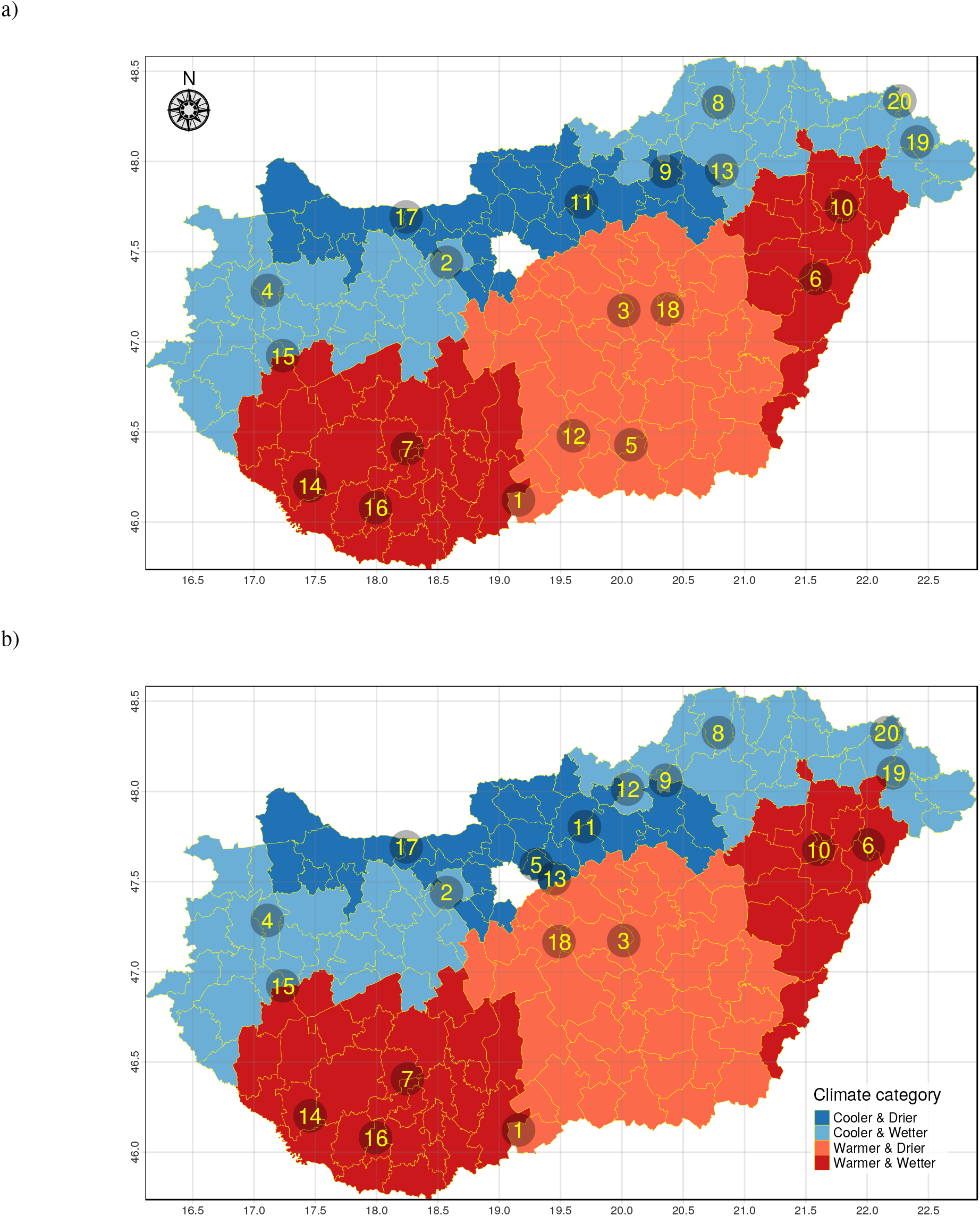
Climate category spatial pattern and sampling points The Hungarian local administrative units (LAU 1) coloured by climatic categories based on growing degree days (GDD) and precipitation of the period 2008-2017. The numbers represent the identification number of the sampled apiaries in March (a) and May (b).

From each involved LAU, one well-managed apiary was selected following personal conversation. Two sample collection campaigns were performed. The first was done between 20/03/2019 and 25/03/2019 (Fig 1/a) and the second in the range of 23/05/2019-01/06/2019 (Fig 1/b), these will be referred to as the sampling periods of March and May, respectively. During the sampling period of March the honey bees were kept on additive feed mostly, with less or no flowers. In May, all apiaries were in blooming acacia (*Robinia pseudoacacia*) forest. For both samplings, the same three-three colonies of the selected apiaries were planed to be involved. At each farm, 20-20 workers of the three colonies were collected and frozen immediately by dry ice. Parallel to the samplings, a survey was conducted to gather metadata on apiaries and their animal health management and status. According to the questionnaires, there was no disease or poisoning in any of the sampled colonies during the study. Since in Hungary mainly Carniolan honey bees (*Apis mellifera carnica*) are in operation, the samples were drawn from that subspecies.

### Sample preparation

The collected samples were prepared for next-generation sequencing (NGS) in the Department of Parasitology and Zoology, University of Veterinary Medicine Budapest. From the deep-frozen workers, 10-10 by colonies were chosen. The bees’ entire gastrointestinal tracts were removed and pooled on the apiary-sampling level. The gut preparation forceps was never used before, and one forceps was applied only for one pool (3×10 guts) processing.

### DNA extraction and metagenomics library preparation

The Quick-DNA Fecal/Soil Microbe Kit from Zymo Research was used for the simple and rapid isolation of inhibitor-free, high-quality host cell and microbial DNA from the bee gut samples. Isolated total metagenome DNA was used for library preparation. In vitro fragment libraries were prepared using the NEBNext Ultra II DNA Library Prep Kit for Illumina. Paired-end fragment reads were generated on an Illumina NextSeq sequencer using TG NextSeq 500/550 High Output Kit v2 (300 cycles). Primary data analysis (base-calling) was carried out with Bbcl2fastq software (v2.17.1.14, Illumina).

### Bioinformatic analysis

After merging the paired-end reads by PEAR^21^ quality-based filtering and trimming was performed by Adapterremoval^22^, using 15 as the quality threshold and only retaining reads longer than 50 bp. The *Apis mellifera* genome (Amel_HAv3.1) sequences as host contaminants were filtered out by Bowtie2^23^ with the very-sensitive-local setting minimising the false positive match level^24^ in further metagenome classification. The remaining reads after deduplication by VSEARCH^25^ were taxonomically classified using Kraken2 (k=35)^26^ with the NCBI non-redundant nucleotide database^27^. Core bacteria was defined as the relative abundance of agglomerated counts on species-level above 0.1% in at least half of the samples. The taxon classification data was managed in R^20^ using functions of package phyloseq^28^ and microbiome^29^.

### Statistical analysis

The within-subject diversity (*α*-diversity) was assessed using the numbers of observed species (richness) and the Inverse Simpson’s Index (evenness). These indices were calculated in 1000 iterations of rarefied OTU tables with a sequencing depth of 6129. The average over the iterations was taken for each apiary. The *α*-diversity expressed by Inverse Simpson’s Index was compared between the conditions using linear models. Comparing the samples collected in March and May, a mixed-effect model was applied to handle the repeated measure by apiary as a random factor.

The between-subject diversity (*β*-diversity) was assessed by Bray-Curtis distance^30^ based on the relative abundances of bacteria species. Using this measure, non-metric multidimensional scaling (NMDS) ordination was applied to visualise the samples’ dissimilarity. To examine statistically whether the bacterial species composition differed by climatic or seasonal conditions PERMANOVA (Permutational Multivariate Analysis of Variance^31^) was performed using vegan package^32^ in R^20^. The abundance differences in core bacteriome between the seasonal or climate conditions were analysed by a negative binomial generalised model of DESeq2 package^33^ in R^20^. This approach was applied following the recommendation of Weiss et al.^34^. None of the compared groups had more than 20 samples, and their average library size ratio was less than 10. Since the apiaries were sampled repeatedly for capturing the seasonal effect, the samples were paired in the model. Considering the multiple comparisons, FDR-adjusted p value (q value) less than 0.10 was considered significant. The statistical tests were two-sided.

## Results

From the twenty apiaries, three-three families were sampled on the first collection period in March. In May there were two apiaries (ID: 6, 13) where one of the original three sampled colonies were not available (e.g. death of the queen, kept at different geographical location). For those apiaries, the pool contained only the two colonies that remained. Eight of the apiaries were migrated (ID: 5, 6, 9, 10, 12, 13, 18, 20), keeping the honey bees in different LAU in March and May. In four apiaries (ID: 5, 9, 12, 13) of these eight, the environmental classification was changed from March to May, because of migration. The apiary ID 5 and ID 12 migrated from a warmer LAU to a cooler one. The apiary ID 9 and ID 12 migrated from an LAU with less precipitation to an LAU with more precipitation. Apiary ID 13 moved from an LAU with more precipitation to an LAU with less precipitation.

The shotgun sequencing generated paired-end read counts of samples ranging between 311,931 and 546,924 with a mean of 413,629. The OTU table created by Kraken2 taxonomic classification contained counts of samples ranging between 11,646 and 114,573 with a mean of 44,280. The minimum, maximum and median read counts of the samples assigned as bacterial species were 6,129, 62,836 and 270,774, respectively.

The numbers of observed species and the Inverse Simpson’s Index *α*-diversity metrics by environmental and seasonal strata are shown in Fig 2. The Inverse Simpson’s Index outliers in the samples collected in March from districts with less and more precipitation are the apiary ID 9 and ID 13, respectively. The apiary ID 12 sampled in March had an outlying high number of observed species too. From the same sampling period among the samples gathered from districts with more precipitation, apiary ID 8 appears as an outlier.

**Figure 2.**
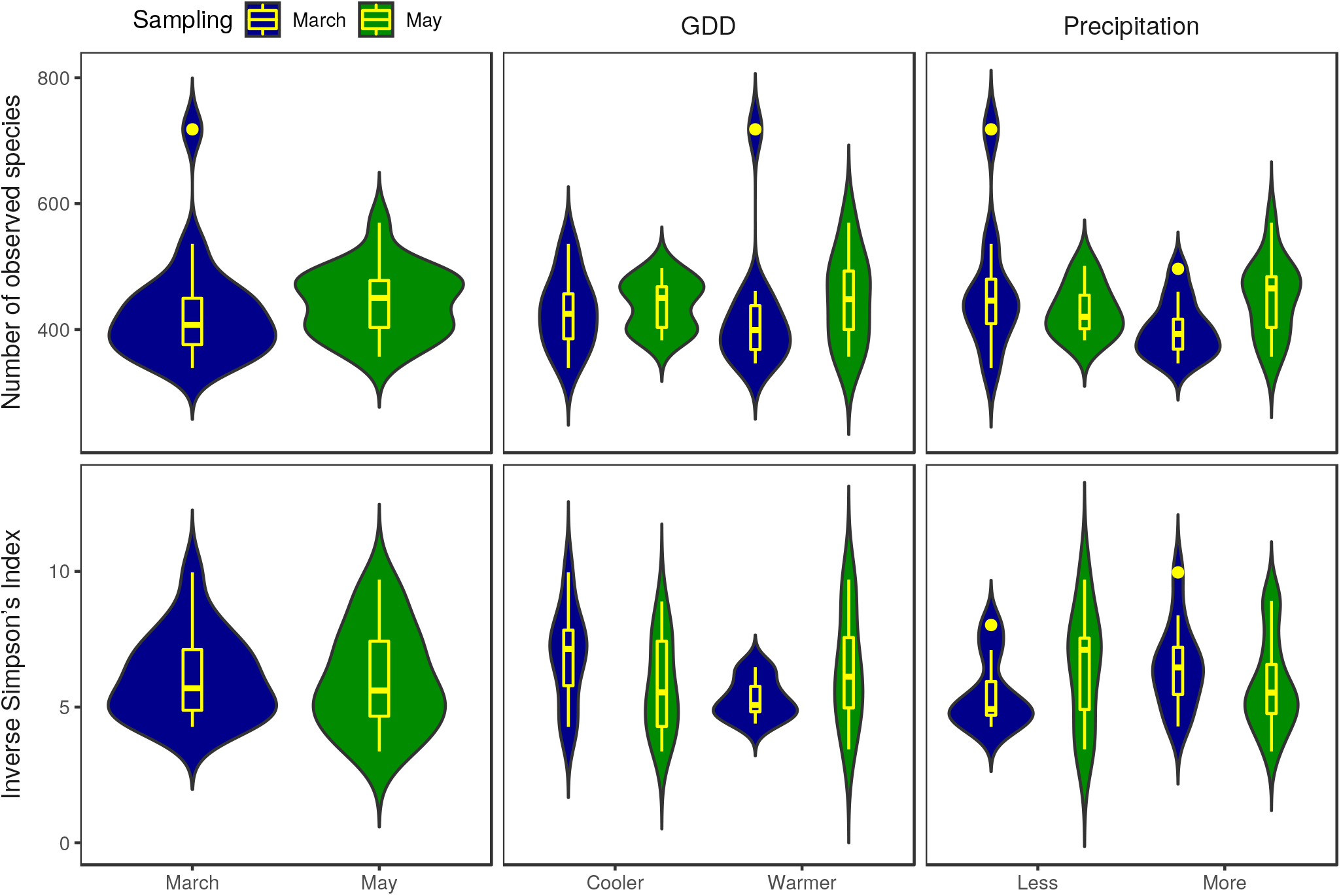
Richness and evenness of honey bee gut bacteriome by sample groups. The numbers of observed species (richness) and the Inverse Simpson’s Index (evenness) as *α*-diversity metrics are presented as a violin and box plot combination. These indices were calculated in 1000 iterations of rarefied OTU tables with a sequencing depth of 6129. The average over the iterations was taken for each apiary. The violin plot shows the probability density, while the box plot marks the outliers, median and the IQR. For Inverse Simpson’s Index, the comparison of samples from cooler and warmer districts collected in March showed significant (p=0.0215) differences only.

In samples from the cooler environment collected in March, the *α*-diversity was significantly (p=0.0215) higher than in samples from warmer districts. There was no significant difference in *α*-diversity between the precipitation categories of samples collected in March (p=0.178). In samples collected in May, there was no significant difference between GDD or precipitation categories (p=0.463 and p=0.456, respectively).

The dissimilarity of the samples’ bacteria species profiles (*β*-diversity) is visualised by NMDS ordination (Fig 3-4), based on Bray-Curtis distance. The ordination stress was 0.144, 0.062 and 0.116 for all samples, samples of March and samples of May, respectively. By PERMANOVA analysis of bacteria species composition significant (p=0.002) difference was found between the samples from March and May. The samples’ bacteriome from March showed similar significant (p=0.02) distance between the cooler and the warmer districts. From the same period, the precipitation levels did not differ significantly (p=0.155). In the samples gathered in May, there was no significant distance between GDD and precipitation categories (p=0.277 and p=0.849, respectively).

**Figure 3.**
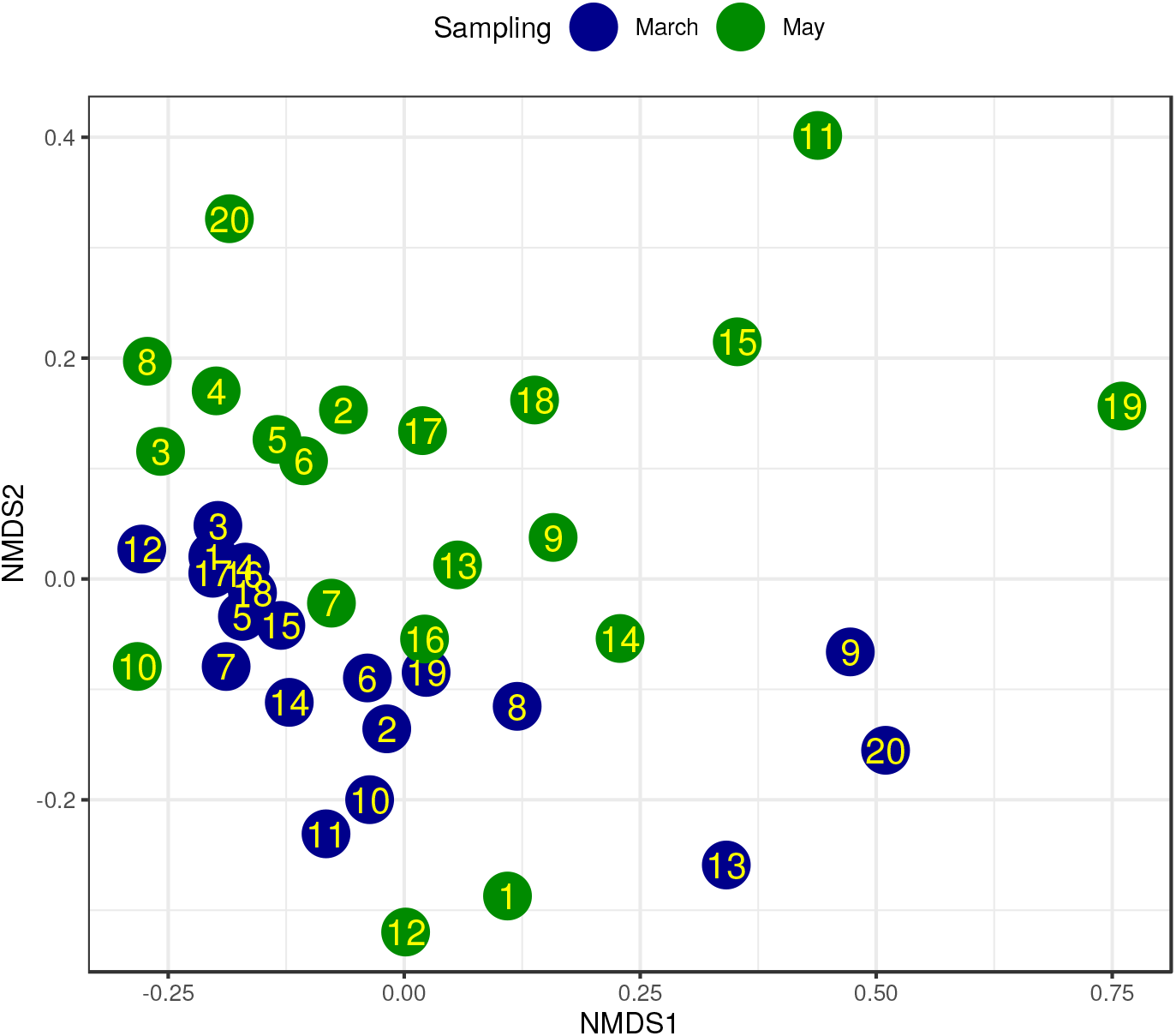
NMDS ordination of bacteriome for sampling March and May. Bray-Curtis dissimilarity was calculated using the species-level abundance of core bacteria. The samples from apiaries (IDs in dots) collected in March (blue) and May (green) are plotted using these dissimilarities. Based on the same measures, PERMANOVA analysis showed significant differences between the sampling time periods (p=0.002, stress=0.144).

**Figure 4.**
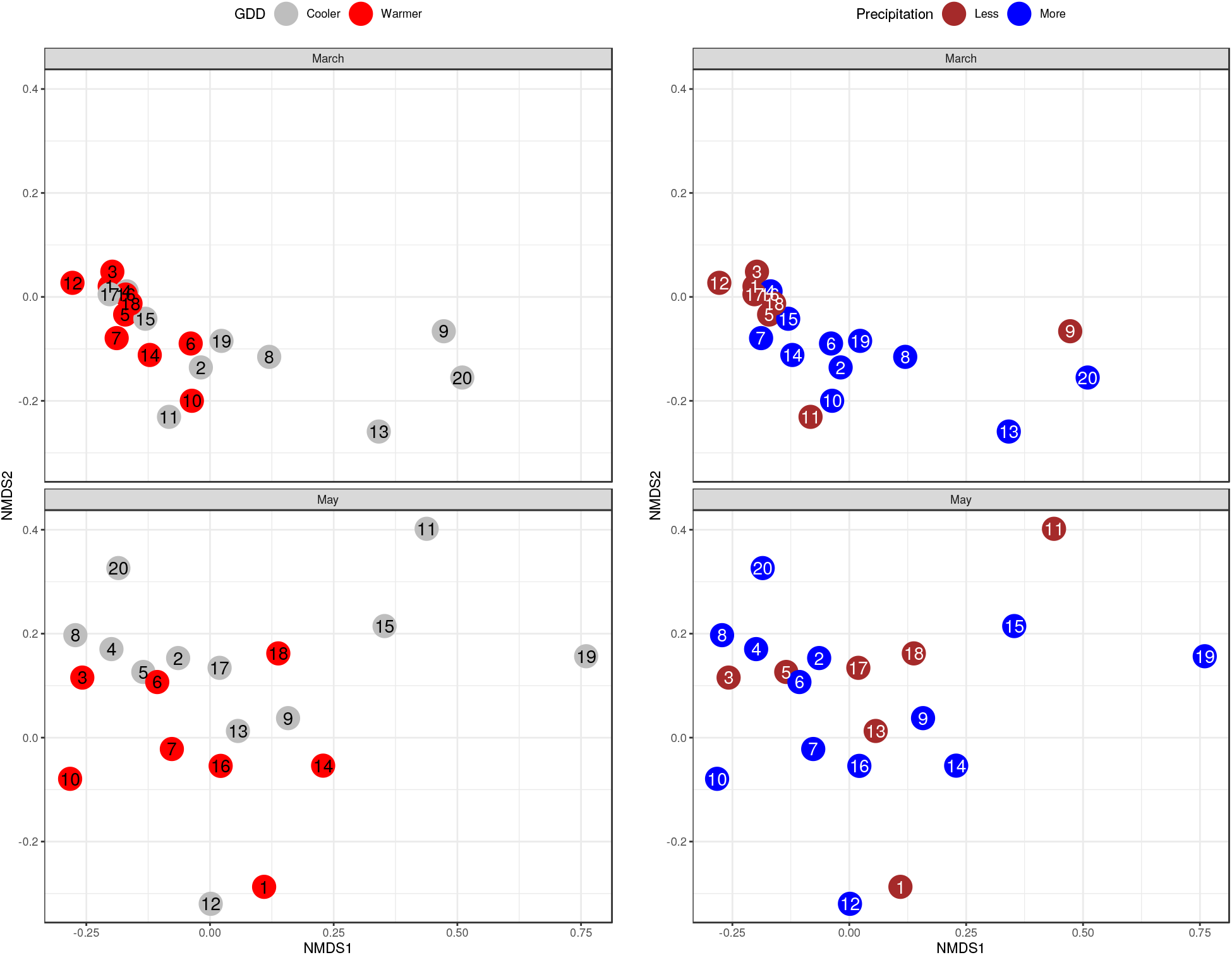
NMDS ordination of bacteriome for environmental condition categories by sampling period. The colours represent the environmental condition categories, and the numbers are corresponding to the apiary IDs. The stress was 0.062 and 0.116 for March and May, respectively. The samples’ bacteriome from March showed significant (p=0.02) distance between the cooler and the warmer districts. From the same period, the precipitation levels did not differ significantly (p=0.155). In the samples gathered in May, there was no significant distance neither between GDD- nor between precipitation categories (p=0.277 and p=0.849, respectively).

The core bacteriome members having relative abundance above 0.1% in at least half of the samples are *Bartonella apis*, *Bifidobacterium asteroides*, *Bifidobacterium coryneforme*, *Bifidobacterium indicum*, *Commensalibacter sp. AMU001*, *Frischella perrara*, *Gilliamella apicola*, *Lactobacillus apis*, *Lactobacillus bombi*, *Lactobacillus helsingborgensis*, *Lactobacillus kullabergensis*, L*actobacillus kunkeei*, *Lactobacillus mellis*, *Lactobacillus sp. wkB8* and *Snodgrassella alvi*. The relative abundances of each apiary’s core bacteriome species are plotted by sampling periods and environmental strata in Fig 5. Table 1 shows the overall and grouped mean and standard deviation of core bacteriome species’ relative abundances.

**Figure 5.**
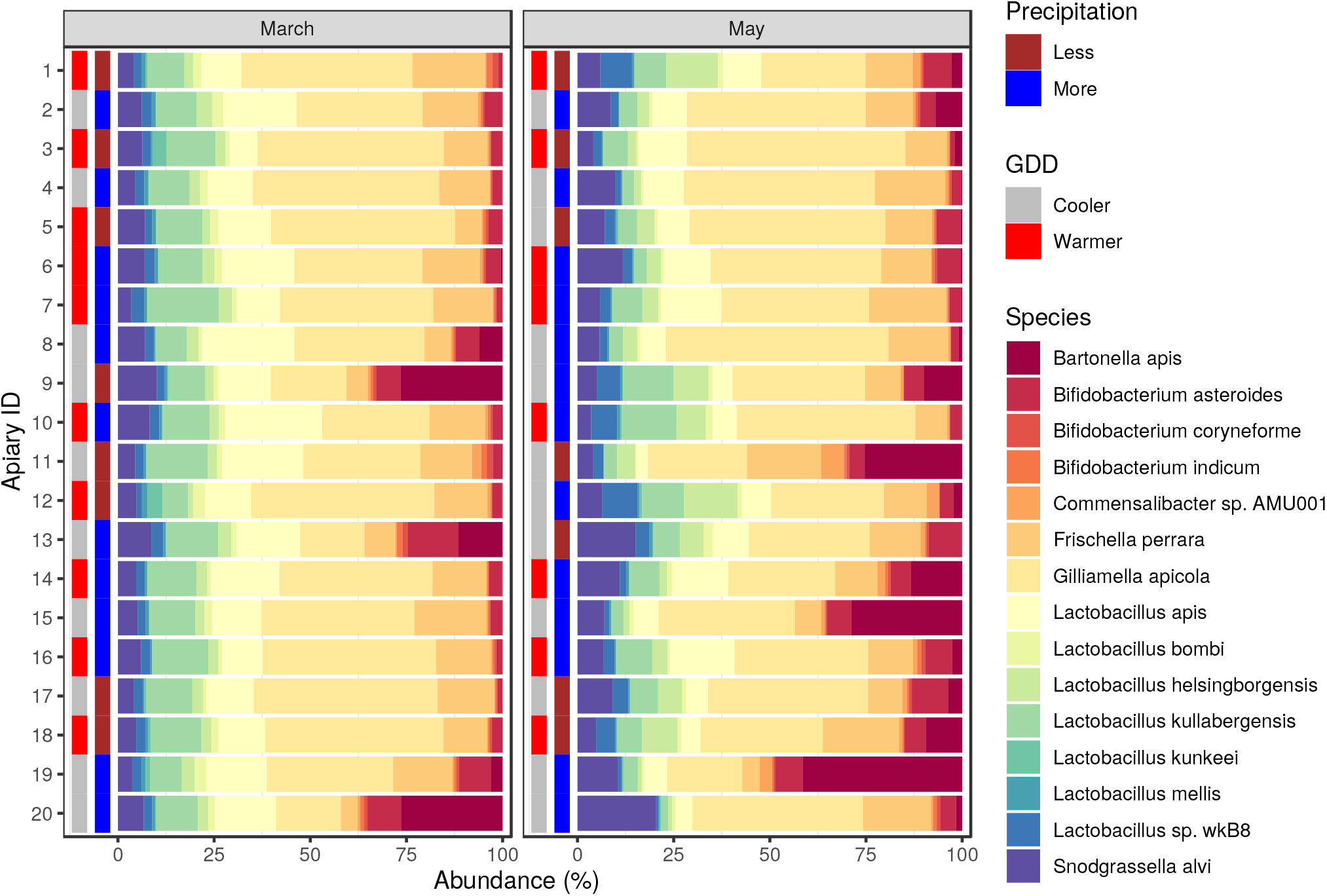
Core bacteriome composition of honey bee gut samples. The relative abundance is plotted for the first (March) and second (May) sampling. Besides the bacteria species of the core bacteriome, the environmental condition categories of sampling places are also marked. The growing degree-day (GDD) categories are coloured by grey and red on the plot’s left sidebar. The next to the right from GDD shows the precipitation categories of apiaries.

**Table 1.** Relative abundances by environmental and seasonal categories.

| Species | All samples<br>Mean (SD) | GDD<br>Cooler | Warmer | Precipitation<br>Less | More |
| --- | --- | --- | --- | --- | --- |
| March |  |  |  |  |  |
| *Bartonella apis* | 3.78 (8.25) | 7.41 (10.7) | 0.14 (0.14) | 3.41 (9.32) | 4.02 (7.89) |
| *Bifidobacterium asteroides* | 4.08 (3.02) | 5.64 (3.62) | 2.52 (0.92) | 2.83 (1.64) | 4.92 (3.49) |
| *Bifidobacterium coryneforme* | 0.64 (0.43) | 0.72 (0.48) | 0.56 (0.38) | 0.73 (0.54) | 0.58 (0.35) |
| *Bifidobacterium indicum* | 0.62 (0.41) | 0.69 (0.44) | 0.55 (0.39) | 0.71 (0.52) | 0.56 (0.33) |
| *Commensalibacter sp. AMU001* | 0.56 (0.49) | 0.66 (0.65) | 0.45 (0.24) | 0.7 (0.73) | 0.46 (0.22) |
| *Frischella perrara* | 12.49 (4.11) | 11.46 (4.85) | 13.52 (3.13) | 12 (4.25) | 12.82 (4.17) |
| *Gilliamella apicola* | 37 (10.63) | 31.92 (11.56) | 42.08 (6.9) | 41.6 (10.67) | 33.93 (9.85) |
| *Lactobacillus apis* | 15.05 (4.73) | 16.23 (4.03) | 13.87 (5.29) | 12.76 (3.93) | 16.57 (4.75) |
| *Lactobacillus bombi* | 1.73 (0.71) | 1.75 (0.78) | 1.71 (0.68) | 1.72 (0.75) | 1.74 (0.72) |
| *Lactobacillus helsingborgensis* | 2.71 (0.61) | 2.91 (0.57) | 2.52 (0.61) | 2.26 (0.49) | 3.01 (0.49) |
| *Lactobacillus kullabergensis* | 11.78 (2.77) | 11.17 (2.42) | 12.4 (3.08) | 11.49 (2.82) | 11.98 (2.84) |
| *Lactobacillus kunkeei* | 0.6 (1.13) | 0.27 (0.35) | 0.94 (1.52) | 1.22 (1.61) | 0.2 (0.3) |
| *Lactobacillus mellis* | 0.75 (0.29) | 0.72 (0.29) | 0.77 (0.31) | 0.77 (0.36) | 0.73 (0.26) |
| *Lactobacillus sp. wkB8* | 2.34 (0.41) | 2.37 (0.28) | 2.3 (0.53) | 2.07 (0.34) | 2.52 (0.37) |
| *Snodgrassella alvi* | 5.88 (1.8) | 6.09 (2.11) | 5.66 (1.5) | 5.74 (2.06) | 5.97 (1.69) |
| May |  |  |  |  |  |
| *Bartonella apis* | 7.6 (11.43) | 10.13 (13.9) | 3.81 (4.87) | 6.19 (8.99) | 8.36 (12.84) |
| *Bifidobacterium asteroides* | 5.12 (2.24) | 5.29 (2.37) | 4.86 (2.16) | 6.11 (2.83) | 4.59 (1.75) |
| *Bifidobacterium coryneforme* | 0.43 (0.29) | 0.42 (0.28) | 0.44 (0.33) | 0.42 (0.2) | 0.43 (0.33) |
| *Bifidobacterium indicum* | 0.44 (0.28) | 0.43 (0.25) | 0.45 (0.34) | 0.41 (0.19) | 0.45 (0.33) |
| *Commensalibacter sp. AMU001* | 1.33 (1.37) | 1.58 (1.66) | 0.96 (0.69) | 1.72 (1.89) | 1.13 (1.02) |
| *Frischella perrara* | 12.79 (4.43) | 12.42 (4.67) | 13.35 (4.28) | 13.79 (4.18) | 12.25 (4.62) |
| *Gilliamella apicola* | 38.71 (10.71) | 38.92 (11.45) | 38.4 (10.25) | 37.91 (12.12) | 39.15 (10.38) |
| *Lactobacillus apis* | 8.74 (3.92) | 6.82 (2.18) | 11.62 (4.29) | 7.67 (3.26) | 9.32 (4.24) |
| *Lactobacillus bombi* | 0.97 (0.5) | 0.95 (0.56) | 1 (0.43) | 1.07 (0.73) | 0.92 (0.34) |
| *Lactobacillus helsingborgensis* | 5.15 (3.81) | 4.74 (3.79) | 5.77 (4) | 6.64 (3.69) | 4.35 (3.76) |
| *Lactobacillus kullabergensis* | 6.52 (3.43) | 5.56 (3.42) | 7.96 (3.08) | 6.26 (1.61) | 6.66 (4.15) |
| *Lactobacillus kunkeei* | 0.18 (0.17) | 0.18 (0.2) | 0.18 (0.14) | 0.12 (0.15) | 0.21 (0.18) |
| *Lactobacillus mellis* | 0.42 (0.19) | 0.41 (0.2) | 0.42 (0.19) | 0.43 (0.27) | 0.4 (0.15) |
| *Lactobacillus sp. wkB8* | 3.44 (2.44) | 3.07 (2.5) | 4.01 (2.4) | 4.06 (2.07) | 3.11 (2.63) |
| *Snodgrassella alvi* | 8.15 (4.16) | 9.09 (4.64) | 6.74 (3.07) | 7.18 (3.91) | 8.67 (4.35) |

Associations between seasonal conditions, climatic condition levels and the abundance of core bacteriome species were examined using negative binomial generalized linear models^33^ (Table 2). The abundance of *Bartonella apis* (FC: 15.41, q<0.00001), *Bifidobacterium asteroides* (FC: 1.61, q=0.0084), *Commensalibacter sp. AMU001* (FC: 2.46, q=0.00001), *Lactobacillus helsingborgensis* (FC: 1.7, q=0.008) and *Snodgrassella alvi* (FC: 1.49, q=0.011) is significantly increased from March to May. In the same comparison the abundance of *Lactobacillus apis* (FC: 0.64, q=0.0066), *Lactobacillus bombi* (FC: 0.64, q=0.0052), *Lactobacillus kullabergensis* (FC: 0.57, q=0.00056) and *Lactobacillus mellis* (FC: 0.64, q=0.0052) was significantly decreased. In the samples collected in March, the abundance of *Lactobacillus kunkeei* (FC: 3.86, q=0.094) was significantly higher in warmer regions than in cooler ones. In the same period, the abundance of *Bartonella apis* (FC: 0.02, q<0.00001) and *Bifidobacterium asteroides* (FC: 0.47, q=0.0027) was significantly lower in warmer LMUs than in cooler ones. In March samples, the abundance of *Bartonella apis* (FC: 11.52, q=0.0066) was significantly higher in districts with more precipitation than in LMUs with less precipitation. The opposite of this was found in the case of *Lactobacillus kunkeei* (FC: 0.13, q=0.00087). In samples collected in May, none of the core bacteriome species showed significant alterations in abundance neither by GDD categories nor by the precipitation levels.

**Table 2.** Abundance alterations of core bacteriome by seasonal and climatic conditions. A negative binomial model estimated the association between species abundance of core bacteriome and sampling seasons, GDD- and precipitation level.

| Samples | Species | Mean counts<sup>†</sup> | Fold change<br>(95% CI) | $q^{\S}$ | Mean counts | Fold change<br>(95% CI) | $q$ |
| --- | --- | --- | --- | --- | --- | --- | --- |
|  |  | GDD<br>Warmer vs. cooler |  |  | Precipitation<br>More vs. less |  |  |
| March | *Bartonella apis* | 1068.87 | 0.02 (0, 0.08) | <0.00001 | *321.94 | 11.52 (2.73, 48.63) | 0.00660 |
|  | *Bifidobacterium asteroides* | 1144.10 | 0.47 (0.31, 0.71) | 0.00274 | 1144.10 | 1.52 (0.94, 2.47) | 0.26310 |
|  | *Bifidobacterium coryneforme* | 180.26 | 0.78 (0.47, 1.3) | 0.55291 | 180.26 | 0.72 (0.43, 1.2) | 0.38749 |
|  | *Bifidobacterium indicum* | 176.76 | 0.81 (0.49, 1.33) | 0.55291 | 176.76 | 0.72 (0.43, 1.19) | 0.38749 |
|  | *Commensalibacter sp. AMU001* | 162.07 | 0.68 (0.37, 1.25) | 0.46158 | 162.07 | 0.58 (0.32, 1.05) | 0.26310 |
|  | *Frischella perrara* | 3662.30 | 1.18 (0.84, 1.65) | 0.55291 | 3662.30 | 0.97 (0.68, 1.37) | 0.84853 |
|  | *Gilliamella apicola* | 11004.31 | 1.29 (0.94, 1.78) | 0.42005 | 11004.31 | 0.75 (0.55, 1.04) | 0.26310 |
|  | *Lactobacillus apis* | 4366.20 | 0.84 (0.65, 1.09) | 0.46158 | 4366.20 | 1.17 (0.89, 1.54) | 0.42531 |
|  | *Lactobacillus bombi* | 495.19 | 1.03 (0.73, 1.44) | 0.86802 | 495.19 | 0.89 (0.63, 1.26) | 0.65673 |
|  | *Lactobacillus helsingborgensis* | 786.53 | 0.87 (0.71, 1.07) | 0.46158 | 786.53 | 1.2 (0.96, 1.5) | 0.26310 |
|  | *Lactobacillus kullabergensis* | 3454.42 | 1.11 (0.87, 1.41) | 0.55291 | 3454.42 | 0.96 (0.74, 1.24) | 0.79727 |
|  | *Lactobacillus kunkei* | 187.49 | 3.86 (1.25, 11.89) | 0.09368 | 187.49 | 0.13 (0.05, 0.36) | 0.00087 |
|  | *Lactobacillus mellis* | 213.99 | 1.13 (0.79, 1.61) | 0.62292 | 213.99 | 0.86 (0.59, 1.25) | 0.56927 |
|  | *Lactobacillus sp. wkB8* | 681.21 | 0.97 (0.8, 1.18) | 0.84023 | 681.21 | 1.1 (0.89, 1.36) | 0.54445 |
|  | *Snodgrassella alvi* | 1706.23 | 0.92 (0.71, 1.2) | 0.62292 | 1706.23 | 0.94 (0.71, 1.24) | 0.77490 |
| May | *Bartonella apis* | 1553.31 | 0.28 (0.08, 1.01) | 0.22572 | 1553.31 | 1.46 (0.36, 5.85) | 0.86244 |
|  | *Bifidobacterium asteroides* | 942.33 | 0.83 (0.56, 1.22) | 0.74756 | 942.33 | 0.9 (0.6, 1.37) | 0.86244 |
|  | *Bifidobacterium coryneforme* | 79.75 | 0.92 (0.49, 1.73) | 0.98269 | 79.75 | 1.13 (0.59, 2.18) | 0.86244 |
|  | *Bifidobacterium indicum* | 81.39 | 0.9 (0.49, 1.63) | 0.98269 | 81.39 | 1.23 (0.66, 2.28) | 0.86244 |
|  | *Commensalibacter sp. AMU001* | 255.76 | 0.51 (0.26, 1) | 0.22572 | *204.88 | 1.33 (0.7, 2.5) | 0.86244 |
|  | *Frischella perrara* | 2522.61 | 1.02 (0.66, 1.56) | 0.98269 | 2522.61 | 0.98 (0.62, 1.54) | 0.92388 |
|  | *Gilliamella apicola* | 7869.43 | 1.01 (0.65, 1.57) | 0.98269 | 7869.43 | 1.09 (0.69, 1.73) | 0.86244 |
|  | *Lactobacillus apis* | 1698.58 | 1.62 (1.07, 2.45) | 0.22572 | 1698.58 | 1.23 (0.77, 1.97) | 0.86244 |
|  | *Lactobacillus bombi* | 183.30 | 1.06 (0.68, 1.66) | 0.98269 | 183.30 | 1.03 (0.65, 1.65) | 0.92388 |
|  | *Lactobacillus helsingborgensis* | 952.28 | 1.2 (0.66, 2.17) | 0.98269 | 952.28 | 0.79 (0.43, 1.45) | 0.86244 |
|  | *Lactobacillus kullabergensis* | 1258.90 | 1.5 (0.93, 2.41) | 0.29310 | 1258.90 | 1.17 (0.7, 1.97) | 0.86244 |
|  | *Lactobacillus kunkei* | 36.06 | 1.08 (0.39, 2.97) | 0.98269 | 36.06 | 1.65 (0.59, 4.57) | 0.86244 |
|  | *Lactobacillus mellis* | 78.82 | 1 (0.65, 1.56) | 0.98269 | 78.82 | 1.12 (0.7, 1.77) | 0.86244 |
|  | *Lactobacillus sp. wkB8* | 648.44 | 1.33 (0.75, 2.36) | 0.74756 | 648.44 | 0.91 (0.49, 1.66) | 0.86244 |
|  | *Snodgrassella alvi* | 1567.32 | 0.66 (0.43, 1.02) | 0.22572 | 1567.32 | 1.43 (0.9, 2.25) | 0.86244 |
| May vs. March |  |  |  |  |  |  |  |
| All | *Bartonella apis* | 1538.45 | 15.41 (6.07, 39.17) | <0.00001 |  |  |  |
|  | *Bifidobacterium asteroides* | 1074.14 | 1.61 (1.16, 2.24) | 0.00837 |  |  |  |
|  | *Bifidobacterium coryneforme* | 118.29 | 0.74 (0.52, 1.05) | 0.12117 |  |  |  |
|  | *Bifidobacterium indicum* | 118.69 | 0.78 (0.55, 1.11) | 0.18588 |  |  |  |
|  | *Commensalibacter sp. AMU001* | 232.29 | 2.46 (1.7, 3.57) | 0.00001 |  |  |  |
|  | *Frischella perrara* | 3046.15 | 1.17 (0.85, 1.61) | 0.34279 |  |  |  |
|  | *Gilliamella apicola* | 9304.64 | 1.22 (0.92, 1.62) | 0.18588 |  |  |  |
|  | *Lactobacillus apis* | 2740.93 | 0.64 (0.48, 0.86) | 0.00656 |  |  |  |
|  | *Lactobacillus bombi* | 309.86 | 0.64 (0.48, 0.84) | 0.00519 |  |  |  |
|  | *Lactobacillus helsingborgensis* | 928.22 | 1.7 (1.19, 2.43) | 0.00800 |  |  |  |
|  | *Lactobacillus kullabergensis* | 2115.13 | 0.57 (0.43, 0.76) | 0.00056 |  |  |  |
|  | *Lactobacillus kunkei* | 99.88 | 0.64 (0.36, 1.14) | 0.16276 |  |  |  |
|  | *Lactobacillus mellis* | 133.69 | 0.64 (0.49, 0.85) | 0.00519 |  |  |  |
|  | *Lactobacillus sp. wkB8* | 680.91 | 1.35 (0.98, 1.87) | 0.10342 |  |  |  |
|  | *Snodgrassella alvi* | 1700.90 | 1.49 (1.12, 1.98) | 0.01099 |  |  |  |

<sup>†</sup> Sequence read counts were normalized by dividing raw counts by DESeq size factors
<sup>§</sup> FDR-adjusted p-value. FDR adjustment was conducted in each pairwise comparison separately
\* Mean counts of two comparisons differ because of outlier replacement

## Discussion

We have examined the bee gut microbiome’s population-level dynamics in different climatic conditions and at different time points in the honey-producing season by determining its *α*- and *β*-diversity. Besides the uniformity of the microbiome’s richness and evenness in almost every examined condition, a significant difference was found between warmer and cooler regions (Figure 2). Considering the differences in the beta-diversity, only the sampling times and the different precipitation levels among the March samples showed significant distinctions (Figures 3 and 4). In accordance with previous findings, where only slight seasonal variations were observed^35, 36^, these results indicate a very conserved, although not unchangeable nature of the honey bee gut microbiome.

Bees possess not only a very constant, but also a rather simple microbiome consisting of only a small subset of phylotypes and associated species, regardless of geographic locations^35–44^. Unsurprisingly, our analysis yielded a similar result to the above-cited articles (Figure 5).

However, on the species level, we found the abundance of several bacterial residents of the honey bee gut to exhibit temporal and environmental differences. On the temporal level, for example, *Snodgrassella alvi*, a microaerophilic Gram-negative bacterium^45^ showed an elevated proportion in the samples collected in May (Figure 5, Table 2). *S. alvi* possibly colonises the gut via an oral-faecal route in the first days of the adult bee’s life^46^. This bacterium forms a biofilm-like layer on the intima of the ileum of the bee gut, along with *Gilliamella apicola*, another characteristic member of the honey bee gut microbiome^38, 40, 41, 47^. *G. apicola* was the most abundant member of the core-bacteriome in our study (Table 1). They do not only have a strong relationship in the biofilm formation, but possibly complement each other in metabolism as well. It was previously shown that *S. alvi*, despite lacking the enzymes of carbohydrate degradation, gain energy from oxidative phosphorylation, which is made possible by utilising the required substrates produced by other microorganisms, such as *G. apicola*^48^. The role of these microbes in the honey bee gut is very diverse, including the defence against different pathogens by biofilm formation^49^. Also, *G. apicola* was shown to exhibit pectinase activity which is important as pectin is a major constituent of the pollen wall^42, 50^. Their relevance is further stressed by the fact that an *S. alvi* strain was found to possess streptomycin resistance genes associated with transposons previously found in *E. coli*^51^. Furthermore, *Snodgrassella* is a candidate microorganism for a bioengineered microbe against bee pathogens^52^.

Besides *S. alvi*, several Lactic Acid Bacteria (LAB), constituting the core-bacteriome in our experiment, showed temporal differences as well (Figure 5, Table 2). They are Gram-positive, catalase-negative, lactic acid fermenting bacteria^53^ and regularly encountered members of the honey bee gut microbiota^37, 43, 54, 55^. Their biofilm-forming capacity was also revealed^54, 56, 57^. LAB species associated with honey bees and bumblebees constitute a highly specialised flora in these insects’ gut which is unique in the insect world^56, 58^. They can metabolise different types of sugars, making them a suitable inhabitant of the intestinal tracts of these species considering that the honey bee diet contains a large amount of carbohydrates. They can also degrade carbohydrates that are indigestible for the host^42, 50, 56^. Nevertheless, it was shown that they even exhibit an ability to metabolise sugars which are toxic to honey bees^42, 50, 56, 59^. Besides the above mentioned nutritional role in the bee gut, they also have a protective function against different pathogenic threats to honey bees^60, 61^. Due to the above-mentioned attributes of the LAB species, it is believed that they are at least partly the reason for the beneficial effects of fresh honey^62^.

Two members of core-bacteriome in this study which showed temporal shifts were *Bartonella apis* and *Commensalibacter sp. AMU001* (both of them were more abundant in the samples collected in May, Figure 5, Table 1). Both of them are non-core members of the honey bee gut microbiome, and members of the *Alphaproteobacteria* class^37, 38, 40, 41, 43, 63, 64^. *Bartonella apis* ferments carbohydrates under microaerophilic conditions^64^. It might also have a role in recycling the end products of honey bee nitrogen metabolism and in the degradation of plant secondary metabolites found in nectar and pollen^65^. *Commensalibacter* genus is a member of the Acetobacteraceae family, which is prevalent in the intestinal tract of many insects, mostly those that consume a sugar-rich diet^66^, for example, butterflies^67^. It was also detected in flowers, possibly originating from visiting insects^68^.

The observed temporal changes in the abundance of the species mentioned above could be due to several reasons. For example, the bees’ foraging behaviour can change by time to match the colonies’ nutritional needs^69^. The relocation of the hives during the season and the seasonal variation of the visited flowers by the foragers^70, 71^ can also account for such changes. Others have also observed temporal changes in the honey bee gut microbiome during the honey-producing season. They found an increased abundance of *Snodgrassella alvi* and *Bartonelly apis* over time and variable changes considering different LAB species. However, it is essential to note that they collected the samples from June to September and only from one location^35^. Kešnerová and colleagues^72^ on the other hand, studied the differences between winter honey bees, foragers and nurses. They observed an increased overall bacterial load with *Snodgrasella*, *Bartonella*, *Commensalibacter* and the two main *Lactobacillus* phylotypes increasing their abundances in winter bees. According to their experiment *Bartonella*, *Commensalibacter* and different *Lactobacillus* species exhibited a decrease in their relative abundances in foragers compared to winter bees in favour of *Snodgrassella* and *Frischella*. Our samples in March could show an intermediate state between full transition from the winter honey bee population to the summer population with the samples in May showing a fully transformed one.

Besides the temporal changes, we also observed differences in the abundance of particular species between various environmental conditions. For example, the Fructophilic Lactic Acid Bacterium (FLAB), *Lactobacillus kunkeei* showed significantly increased abundance in the warmer and drier regions in samples collected in May (Table 2).

FLAB is a distinct and special group of LAB, whose main characteristics are the preference of fructose over glucose and that they can metabolise glucose only in the presence of external electron acceptors (e.g. pyruvate or oxygen)^73, 74^. Considering the honey bee’s fructose-rich diet and the high sugar tolerance of FLAB species, it makes them suitable for this environment^74, 75^.

*L. kunkeei* is the most important FLAB species found in the honey bee intestinal tract and is most abundantly found in the honey stomach^36, 55, 57, 73–76^. Although, *L. kunkeei* is usually one of the most abundant members of the honey bee intestinal tract^41^, in this case, we found *Lactobacillus apis* and *Lactobacillus kullabergensis* to be (Figure 5, Table 1). However, it is an important member of the bee microbiome and *L. kunkeei* can be found in several bee products, such as honey or bee bread^55, 75, 77^. It can also be found in flowers and fruits^73, 78^ and can thus get into the products made from them. For example, *L. kunkeei* was originally isolated from wine^79^. This species’ previous gene sequence analysis revealed a major gene loss^80^ accompanied with an enrichment of genes involved in fructose metabolism and lipid- and amino acid synthesis^80–82^. The genetic composition of this species indicates its high adaptation to sugar (mainly fructose) rich environments. Besides its highly adapted nature to this environment, *L. kunkeei* was shown to have developed different techniques to protect its ecological niche. Phenotypical and genotypical experiments demonstrated the ability of *L. kunkeei* to form biofilms, which is a major advantage in antimicrobial activity. This potential makes it a good candidate probiotic for either humans or insects^81, 83, 84^. These attributes make *L. kunkeei* a protective barrier against environmental pathogens of honey bees. It was found to be effective against different pathogens, including *Nosema ceraneae*, *Paenibacillus larvae* and *Melissococcus plutonius*^75, 85^. Also, the MP2 strain of *L. kunkeei* was shown to possess genes for isoprenoid synthesis, indicating its possible role in honey bee nutrition^81^. Its effect was even determined in bee bread maturation^77^.

Changes in abundance for different environmental conditions were also found in the case *Bartonella apis* (more abundant in colder regions with higher precipitation, Table 2) and *Bifidobacterium asteroides* (more abundant in colder regions, Table 2).

The differences observed under various environmental conditions can be explained by the fact that these effects, such as, temperature or exposure to sunlight can affect the nectar production and nectar composition of plant species^86–88^. Nevertheless, it is essential to note the differences in the plant species composition of each ecological region. The different flowers that honey bees are foraging can affect, for example, the honey stomach microbiota^55^. It was also found that the microbiome differed in foragers kept feeding on flowering oilseed rape compared to those foraging other pollen and nectar sources. However, these effects could originate from the different neonicotinoid contamination of plants^39^.

*Frischella perrara* is also commonly found in the gut of honey bees^38, 40, 41, 89^, and it was found to be a member of the core-bacteriome in our study as well. Although it was present here, we could not reveal any variation in its abundance neither temporally, nor between different environmental conditions (Figure 5, Table 1, Table 2). *F. perrara* is a close relative of *G. apicola*, and has a similar metabolism^89^. It is usually found in the ileum of the bee gut^90^ and was shown to share a strong relationship with the honey bee immune system^91^.

Although consisting only of a few species, the honey bee microbiome possesses several functions affecting its host’s health. It has a role in the degradation of different polysaccharides originating from the bees’ diet, for example, from pollen walls. These are often undegradable for bees^42, 92^. They might also have a role in recycling the nitrogen waste materials of honey bee nitrogen metabolism^92^ and can metabolise potentially toxic sugars for bees^42^. Besides these metabolic functions, the gut microbiome has a positive impact on the host immune system^93^, and it protects the bees from pathogens^93, 94^.

Considering the honey bee gut microbiome’s diverse role, everything affecting it can potentially threaten bee health. For example, it was shown that the widely used herbicide, glyphosate, impacts the microbiota^9, 10^ and other stressors can induce changes in its composition as well^93, 95–97^. However, to understand and evaluate the potential impact of these effects on honey bees, we need to know the microbiota’s normal composition and function.

## Conclusion

Based on our results, one may conclude that the composition of healthy core bacteriomes in honey bees varies depending on the climatic and seasonal conditions. This is probably since climatic characteristics and vegetation states determine the availability and nutrient content of flowering plants. The results of our study prove that in order to gain a thorough understanding of a microbiome’s natural diversity, we need to obtain the necessary information from extreme ranges of the host’s healthy state.

## Acknowledgements

In memory of Rajnald András Köveshegyi OCist. We would like to say thanks to the beekeepers for giving us their indispensable help. The project is supported by the European Union and co-financed by the European Social Fund (No. EFOP-3.6.3-VEKOP-16-2017-00005). It has also received funding from the European Union’s Horizon 2020 research and innovation program under Grant Agreement No. 874735 (VEO). GM received support from the Hungarian Academy of Sciences through the Lendület-Programme (LP2020-5/2020).

## Author contributions statement

NS takes responsibility for the integrity of the data and the accuracy of the data analysis. LB, LM, NS and RF conceived the concept of the study. GM, LB, LM, NS and RF performed sample collection and procedures. MP and NS participated in the bioinformatic and statistical analysis. MP and NS participated in the drafting of the manuscript. DT, GM, LB, LM, MP, NS, and RF carried out the manuscript’s critical revision for important intellectual content. All authors read and approved the final manuscript.

## Additional information

### Availability of data and material

The short read data of samples are publicly available and accessible through the PR-JNA685398 from the NCBI Sequence Read Archive (SRA). Reviewer link: https://dataview.ncbi.nlm.nih.gov/object/PRJNA685398?reviewer=3if5rmnvevgdq3qskhos2dlj4r

### Ethics approval and consent to participate

Not applicable.

### Consent for publication

Not applicable.

## References

1. Ványi, G. Á., Csapó, Z. & Kárpáti, L. Externality effects of honey production. Appl. Stud. Agribusiness Commer. 6, 63–67 (2012).

2. Hristov, P., Neov, B., Shumkova, R. & Palova, N. Significance of apoidea as main pollinators. ecological and economic impact and implications for human nutrition. Diversity 12, 280 (2020).

3. Patel, V., Pauli, N., Biggs, E., Barbour, L. & Boruff, B. Why bees are critical for achieving sustainable development. Ambio 50, 49–59 (2021).

4. Samarghandian, S., Farkhondeh, T. & Samini, F. Honey and health: A review of recent clinical research (2017).

5. Oldroyd, B. P. What’s killing american honey bees? PLoS Biol 5, e168 (2007).

6. Barbosa, W. F., Smagghe, G. & Guedes, R. N. C. Pesticides and reduced-risk insecticides, native bees and pantropical stingless bees: pitfalls and perspectives. Pest Manag. Sci. 71, 1049–1053 (2015).

7. Morawetz, L. et al. Health status of honey bee colonies (apis mellifera) and disease-related risk factors for colony losses in austria. PloS one 14, e0219293 (2019).

8. Potts, S. G. et al. Global pollinator declines: trends, impacts and drivers. Trends ecology & evolution 25, 345–353 (2010).

9. Dai, P. et al. The herbicide glyphosate negatively affects midgut bacterial communities and survival of honey bee during larvae reared in vitro. J. Agric. Food Chem. 66, 7786–7793 (2018).

10. Motta, E. V., Raymann, K. & Moran, N. A. Glyphosate perturbs the gut microbiota of honey bees. Proc. Natl. Acad. Sci. United States Am. 115, 10305–10310 (2018).

11. Paris, L. et al. Honeybee gut microbiota dysbiosis in pesticide/parasite co-exposures is mainly induced by nosema ceranae. J. invertebrate pathology 172, 107348 (2020).

12. Moran, N. A., Hansen, A. K., Powell, J. E. & Sabree, Z. L. Distinctive gut microbiota of honey bees assessed using deep sampling from individual worker bees. PLoS ONE 7(2012).

13. Anderson, K. E. et al. The queen’s gut refines with age: longevity phenotypes in a social insect model. Microbiome 6, 108 (2018).

14. Regan, T. et al. Characterisation of the UK honey bee (Apis mellifera) metagenome. bioRxiv 293647 (2018).

15. Petersen, J. D., Reiners, S. & Nault, B. A. Pollination Services Provided by Bees in Pumpkin Fields Supplemented with Either *Apis mellifera* or *Bombus impatiens* or Not Supplemented. PLoS ONE 8, e69819 (2013).

16. Gilley, D. C., Courtright, T. J. & Thom, C. Phenology of Honey Bee Swarm Departure in New Jersey, United States. Environ. Entomol. 47, 603–608 (2018).

17. Dee, D. P. et al. The ERA-Interim reanalysis: configuration and performance of the data assimilation system. Q. J. Royal Meteorol. Soc. 137, 553–597 (2011).

18. Stevens, D. L. & Olsen, A. R. Spatially balanced sampling of natural resources. J. Am. Stat. Assoc. 99, 262–278 (2004).

19. Kincaid, T. M., Olsen, A. R. & Weber, M. H. spsurvey: Spatial Survey Design and Analysis (2019). R package version 4.1.0.

20. R Core Team. R: A Language and Environment for Statistical Computing. R Foundation for Statistical Computing, Vienna, Austria (2020).

21. Zhang, J., Kobert, K., Flouri, T. & Stamatakis, A. Pear: a fast and accurate illumina paired-end read merger. Bioinformatics 30, 614–620 (2013).

22. Schubert, M., Lindgreen, S. & Orlando, L. AdapterRemoval v2: rapid adapter trimming, identification, and read merging. BMC Res. Notes 9, 88 (2016).

23. Langmead, B. & Salzberg, S. L. Fast gapped-read alignment with Bowtie 2. Nat. Methods 9, 357–359 (2012).

24. Czajkowski, M. D., Vance, D. P., Frese, S. A. & Casaburi, G. GenCoF: a graphical user interface to rapidly remove human genome contaminants from metagenomic datasets. Bioinformatics (2018).

25. Rognes, T., Flouri, T., Nichols, B., Quince, C. & Mahé, F. VSEARCH: a versatile open source tool for metagenomics. PeerJ 4, e2584 (2016).

26. Wood, D. E., Lu, J. & Langmead, B. Improved metagenomic analysis with kraken 2. Genome biology 20, 1–13 (2019).

27. Pruitt, K. D., Tatusova, T. & Maglott, D. R. NCBI Reference Sequence (RefSeq): a curated non-redundant sequence database of genomes, transcripts and proteins. Nucleic acids research 33, D501–4 (2005).

28. McMurdie, P. J. & Holmes, S. phyloseq: An r package for reproducible interactive analysis and graphics of microbiome census data. PLOS ONE 8, 1–11 (2013).

29. Lahti, L. & Shetty, S. microbiome r package (2012-2019).

30. Bray, J. R. & Curtis, J. T. An Ordination of the Upland Forest Communities of Southern Wisconsin. Ecol. Monogr. 27, 325–349 (1957).

31. Anderson, M. J. A new method for non-parametric multivariate analysis of variance. Austral ecology 26, 32–46 (2001).

32. Oksanen, J. et al. vegan: Community Ecology Package (2019). R package version 2.5-6.

33. Love, M. I., Huber, W. & Anders, S. Moderated estimation of fold change and dispersion for RNA-seq data with DESeq2. Genome Biol. 15, 550 (2014).

34. Weiss, S. et al. Normalization and microbial differential abundance strategies depend upon data characteristics. Microbiome 5, 27 (2017).

35. Subotic, S. et al. Honey bee microbiome associated with different hive and sample types over a honey production season. PloS one 14(2019).

36. Corby-Harris, V., Maes, P. & Anderson, K. E. The bacterial communities associated with honey bee (apis mellifera) foragers. PloS one 9(2014).

37. Ellegaard, K. M. & Engel, P. Genomic diversity landscape of the honey bee gut microbiota. Nat. communications 10, 1–13 (2019).

38. Jones, J. C. et al. The gut microbiome is associated with behavioural task in honey bees. Insectes sociaux 65, 419–429 (2018).

39. Jones, J. C. et al. Gut microbiota composition is associated with environmental landscape in honey bees. Ecol. evolution 8, 441–451 (2018).

40. Kwong, W. K. & Moran, N. A. Gut microbial communities of social bees. Nat. Rev. Microbiol. 14, 374 (2016).

41. Anderson, K. E. et al. Microbial ecology of the hive and pollination landscape: bacterial associates from floral nectar, the alimentary tract and stored food of honey bees (apis mellifera). PloS one 8(2013).

42. Engel, P., Martinson, V. G. & Moran, N. A. Functional diversity within the simple gut microbiota of the honey bee. Proc. Natl. Acad. Sci. 109, 11002–11007 (2012).

43. Martinson, V. G. et al. A simple and distinctive microbiota associated with honey bees and bumble bees. Mol. Ecol. 20, 619–628 (2011).

44. Cox-Foster, D. L. et al. A metagenomic survey of microbes in honey bee colony collapse disorder. Science 318, 283–287 (2007).

45. Kwong, W. K. & Moran, N. A. Cultivation and characterization of the gut symbionts of honey bees and bumble bees: description of snodgrassella alvi gen. nov., sp. nov., a member of the family neisseriaceae of the betaproteobacteria, and gilliamella apicola gen. nov., sp. nov., a member of orbaceae fam. nov., orbales ord. nov., a sister taxon to the order ‘enterobacteriales’ of the gammaproteobacteria. Int. journal systematic evolutionary microbiology 63, 2008–2018 (2013).

46. Powell, J. E., Martinson, V. G., Urban-Mead, K. & Moran, N. A. Routes of acquisition of the gut microbiota of the honey bee apis mellifera. Appl. Environ. Microbiol. 80, 7378–7387 (2014).

47. Martinson, V. G., Moy, J. & Moran, N. A. Establishment of characteristic gut bacteria during development of the honeybee worker. Appl. Environ. Microbiol. 78, 2830–2840 (2012).

48. Kwong, W. K., Engel, P., Koch, H. & Moran, N. A. Genomics and host specialization of honey bee and bumble bee gut symbionts. Proc. Natl. Acad. Sci. 111, 11509–11514 (2014).

49. Cariveau, D. P., Powell, J. E., Koch, H., Winfree, R. & Moran, N. A. Variation in gut microbial communities and its association with pathogen infection in wild bumble bees (bombus). The ISME journal 8, 2369–2379 (2014).

50. Lee, F. J., Miller, K. I., McKinlay, J. B. & Newton, I. L. Differential carbohydrate utilization and organic acid production by honey bee symbionts. FEMS microbiology ecology 94, fiy113 (2018).

51. Ludvigsen, J., Amdam, G. V., Rudi, K. & L’Abée-Lund, T. M. Detection and characterization of streptomycin resistance (stra-strb) in a honeybee gut symbiont (snodgrassella alvi) and the associated risk of antibiotic resistance transfer. Microb. ecology 76, 588–591 (2018).

52. Leonard, S. P. et al. Engineered symbionts activate honey bee immunity and limit pathogens. Science 367, 573–576 (2020).

53. Felis, G. E. & Dellaglio, F. Taxonomy of lactobacilli and bifidobacteria. Curr. issues intestinal microbiology 8, 44 (2007).

54. Audisio, M. C., Torres, M. J., Sabaté, D. C., Ibarguren, C. & Apella, M. C. Properties of different lactic acid bacteria isolated from apis mellifera l. bee-gut. Microbiol. research 166, 1–13 (2011).

55. Olofsson, T. C. & Vásquez, A. Detection and identification of a novel lactic acid bacterial flora within the honey stomach of the honeybee apis mellifera. Curr. microbiology 57, 356–363 (2008).

56. Ellegaard, K. M. et al. Extensive intra-phylotype diversity in lactobacilli and bifidobacteria from the honeybee gut. BMC genomics 16, 284 (2015).

57. Vásquez, A. et al. Symbionts as major modulators of insect health: lactic acid bacteria and honeybees. PloS one 7(2012).

58. McFrederick, Q. S. et al. Specificity between lactobacilli and hymenopteran hosts is the exception rather than the rule. Appl. Environ. Microbiol. 79, 1803–1812 (2013).

59. Sols, A., Cadenas, E. & Alvarado, F. Enzymatic basis of mannose toxicity in honey bees. Science 131, 297–298 (1960).

60. Baffoni, L. et al. Effect of dietary supplementation of bifidobacterium and lactobacillus strains in apis mellifera l. against nosema ceranae. Benef. microbes 7, 45–51 (2016).

61. Killer, J., Dubná, S., Sedláček, I. & Švec, P. Lactobacillus apis sp. nov., from the stomach of honeybees (apis mellifera), having an in vitro inhibitory effect on the causative agents of american and european foulbrood. Int. journal systematic evolutionary microbiology 64, 152–157 (2014).

62. Olofsson, T. C. et al. Lactic acid bacterial symbionts in honeybees–an unknown key to honey’s antimicrobial and therapeutic activities. Int. wound journal 13, 668–679 (2016).

63. Siozios, S., Moran, J., Chege, M., Hurst, G. D. & Paredes, J. C. Complete reference genome assembly for commensalibacter sp. strain amu001, an acetic acid bacterium isolated from the gut of honey bees. Microbiol Resour Announc. 8, e01459–18 (2019).

64. Kešnerová, L., Moritz, R. & Engel, P. Bartonella apis sp. nov., a honey bee gut symbiont of the class alphaproteobacteria. Int. journal systematic evolutionary microbiology 66, 414–421 (2016).

65. Segers, F. H., Kešnerová, L., Kosoy, M. & Engel, P. Genomic changes associated with the evolutionary transition of an insect gut symbiont into a blood-borne pathogen. The ISME journal 11, 1232–1244 (2017).

66. Crotti, E. et al. Acetic acid bacteria, newly emerging symbionts of insects. Appl. Environ. Microbiol. 76, 6963–6970 (2010).

67. Hammer, T. J., McMillan, W. O. & Fierer, N. Metamorphosis of a butterfly-associated bacterial community. PloS one 9 (2014).

68. Morris, M. M., Frixione, N. J., Burkert, A. C., Dinsdale, E. A. & Vannette, R. L. Microbial abundance, composition, and function in nectar are shaped by flower visitor identity. FEMS Microbiol. Ecol. (2020).

69. Hendriksma, H. P. & Shafir, S. Honey bee foragers balance colony nutritional deficiencies. Behav. Ecol. Sociobiol. 70, 509–517 (2016).

70. Bertrand, C. et al. Seasonal shifts and complementary use of pollen sources by two bees, a lacewing and a ladybeetle species in european agricultural landscapes. J. Appl. Ecol. 56, 2431–2442 (2019).

71. Requier, F. et al. Honey bee diet in intensive farmland habitats reveals an unexpectedly high flower richness and a major role of weeds. Ecol. Appl. 25, 881–890 (2015).

72. Kešnerová, L. et al. Gut microbiota structure differs between honeybees in winter and summer. The ISME journal 14, 801–814 (2020).

73. Neveling, D. P., Endo, A. & Dicks, L. M. Fructophilic lactobacillus kunkeei and lactobacillus brevis isolated from fresh flowers, bees and bee-hives. Curr. microbiology 65, 507–515 (2012).

74. Endo, A., Futagawa-Endo, Y. & Dicks, L. M. Isolation and characterization of fructophilic lactic acid bacteria from fructose-rich niches. Syst. Appl. Microbiol. 32, 593–600 (2009).

75. Endo, A. & Salminen, S. Honeybees and beehives are rich sources for fructophilic lactic acid bacteria. Syst. Appl. Microbiol. 36, 444–448 (2013).

76. Endo, A. et al. Characterization and emended description of lactobacillus kunkeei as a fructophilic lactic acid bacterium. Int. journal systematic evolutionary microbiology 62, 500–504 (2012).

77. Di Cagno, R., Filannino, P., Cantatore, V. & Gobbetti, M. Novel solid-state fermentation of bee-collected pollen emulating the natural fermentation process of bee bread. Food microbiology 82, 218–230 (2019).

78. Sakandar, H. A., Kubow, S. & Sadiq, F. A. Isolation and in-vitro probiotic characterization of fructophilic lactic acid bacteria from chinese fruits and flowers. LWT 104, 70–75 (2019).

79. Edwards, C., Haag, K., Collins, M., Hutson, R. & Huang, Y. Lactobacillus kunkeei sp. nov.: a spoilage organism associated with grape juice fermentations. J. applied microbiology 84, 698–702 (1998).

80. Tamarit, D. et al. Functionally structured genomes in lactobacillus kunkeei colonizing the honey crop and food products of honeybees and stingless bees. Genome Biol. Evol. 7, 1455–1473 (2015).

81. Asenjo, F. et al. Genome sequencing and analysis of the first complete genome of lactobacillus kunkeei strain mp2, an apis mellifera gut isolate. PeerJ 4, e1950 (2016).

82. Maeno, S. et al. Genomic characterization of a fructophilic bee symbiont lactobacillus kunkeei reveals its niche-specific adaptation. Syst. applied microbiology 39, 516–526 (2016).

83. Berríos, P. et al. Inhibitory effect of biofilm-forming lactobacillus kunkeei strains against virulent pseudomonas aeruginosa in vitro and in honeycomb moth (galleria mellonella) infection model. Benef. microbes 9, 257–268 (2018).

84. Djukic, M. et al. High quality draft genome of lactobacillus kunkeei efb6, isolated from a german european foulbrood outbreak of honeybees. Standards genomic sciences 10, 16 (2015).

85. Arredondo, D. et al. Lactobacillus kunkeei strains decreased the infection by honey bee pathogens paenibacillus larvae and nosema ceranae. Benef. Microbes 9, 279–290 (2018).

86. Takkis, K., Tscheulin, T. & Petanidou, T. Differential effects of climate warming on the nectar secretion of early-and late-flowering mediterranean plants. Front. plant science 9, 874 (2018).

87. Takkis, K., Tscheulin, T., Tsalkatis, P. & Petanidou, T. Climate change reduces nectar secretion in two common mediterranean plants. AoB Plants 7(2015).

88. Nocentini, D., Pacini, E., Guarnieri, M., Martelli, D. & Nepi, M. Intrapopulation heterogeneity in floral nectar attributes and foraging insects of an ecotonal mediterranean species. Plant ecology 214, 799–809 (2013).

89. Engel, P., Kwong, W. K. & Moran, N. A. Frischella perrara gen. nov., sp. nov., a gammaproteobacterium isolated from the gut of the honeybee, apis mellifera. Int. journal systematic evolutionary microbiology 63, 3646–3651 (2013).

90. Engel, P., Bartlett, K. D. & Moran, N. A. The bacterium frischella perrara causes scab formation in the gut of its honeybee host. MBio 6, e00193–15 (2015).

91. Emery, O., Schmidt, K. & Engel, P. Immune system stimulation by the gut symbiont frischella perrara in the honey bee (apis mellifera). Mol. ecology 26, 2576–2590 (2017).

92. Zheng, H. et al. Division of labor in honey bee gut microbiota for plant polysaccharide digestion. Proc. Natl. Acad. Sci. 116, 25909–25916 (2019).

93. Raymann, K. & Moran, N. A. The role of the gut microbiome in health and disease of adult honey bee workers. Curr. opinion insect science 26, 97–104 (2018).

94. Bonilla-Rosso, G. & Engel, P. Functional roles and metabolic niches in the honey bee gut microbiota. Curr. opinion microbiology 43, 69–76 (2018).

95. Rouzé, R., Moné, A., Delbac, F., Belzunces, L. & Blot, N. The honeybee gut microbiota is altered after chronic exposure to different families of insecticides and infection by nosema ceranae. Microbes environments ME18169 (2019).

96. Li, J. et al. Pollen reverses decreased lifespan, altered nutritional metabolism and suppressed immunity in honey bees (apis mellifera) treated with antibiotics. J. Exp. Biol. 222(2019).

97. Raymann, K., Shaffer, Z. & Moran, N. A. Antibiotic exposure perturbs the gut microbiota and elevates mortality in honeybees. PLoS biology 15(2017).

